# Tracking a Serial Killer: Integrating Phylogenetic Relationships, Epidemiology, and Geography for Two Invasive Meningococcal Disease Outbreaks

**DOI:** 10.1101/387597

**Authors:** Ifeoma Ezeoke, Madeline R. Galac, Ying Lin, Alvin T. Liem, Pierce A. Roth, Andrew Kilianski, Henry S. Gibbons, Danielle Bloch, John Kornblum, Paula Del Rosso, Daniel A. Janies, Don Weiss

## Abstract

**Background:** While overall rates of meningococcal disease have been declining in the United States for the past several decades, New York City (NYC) has experienced two serogroup C meningococcal disease outbreaks in 2005-2006 and in 2010-2013. The outbreaks were centered within drug use and sexual networks, were difficult to control, and required vaccine campaigns.

**Methods:** Whole Genome Sequencing (WGS) was used to analyze preserved meningococcal isolates collected before and during the two outbreaks. We integrated and analyzed epidemiologic, geographic, and genomic data to better understand transmission networks among patients. Betweenness centrality was used as a metric to understand the most important geographic nodes in the transmission networks. Comparative genomics was used to identify genes associated with the outbreaks.

**Results:** *Neisseria meningitidis* serogroup C (ST11/ET-37) was responsible for both outbreaks with each outbreak having distinct phylogenetic clusters. WGS did identify some misclassifications of isolates that were more distant from the rest of the outbreak, as well as those that should have been included based on high genomic similarity. Genomes for the second outbreak were more similar than the first and no mutation was found to either be unique or specific to either outbreak lineage. Betweenness centrality as applied to transmission networks based on phylogenetic analysis demonstrated that the outbreaks were transmitted within focal communities in NYC with few transmission events to other locations.

**Conclusions:** *Neisseria meningitidis* is an ever changing pathogen and comparative genomic analyses can help elucidate how it spreads geographically to facilitate targeted interventions to interrupt transmission.

## BACKGROUND

The incidence of meningococcal disease in the United States has declined steadily since 1996 [1]. Against this backdrop, New York City (NYC) experienced 2 outbreaks of invasive meningococcal disease (IMD) caused by serogroup C *Neisseria meningitidis* (MenC) that prompted vaccination interventions [2, 3]. The first NYC MenC outbreak (Ob1) occurred in the years 2005-2006 and affected current and former drug users and their close contacts [2]. The second community outbreak of IMD (Ob2) began four years later and affected 22 men who have sex with men (MSM) [3]. Previous outbreaks of IMD among MSM were seen in the early 2000s in Toronto [4] and Chicago [5] and after the NYC outbreak, clusters of IMD in MSM were recognized in Southern California [6], Chicago [7], Germany [8], France [9], and Spain [10].

*N. meningitidis* is found in the nasopharynx in up to 35% of healthy adults [11] and it is this colonization state that is believed to be responsible for IMD transmission and outbreaks. Secondary IMD cases are rare and outbreaks occur in either organizational settings, such as universities, or in members of discreet community groups, such as MSM. Pulsed-field gel electrophoresis (PFGE) has been used in NYC to detect links among IMD cases not identified by case investigations [12]. However, PFGE may misclassify due to a lack of discriminatory power [13]. Additionally, PFGE does not provide details on mutations in the *N. meningitidis* genome that can be used to track its spread in space and time.

Advances in whole genome sequencing (WGS) technology have allowed epidemiologists to uncover disease transmission patterns through phylogenetic analyses. WGS has successfully been used to inform outbreaks caused by tuberculosis [14], cholera [15], *Klebsiella* [16], *Legionella* [17], and *Listeria* [18]. Also, WGS has led to two recent reports suggesting that *N. meningitidis* has adapted to become a urogenital pathogen. Taha et al. sequenced *N. meningitidis* isolates from the 2012-13 MSM outbreak in Germany and France and several antecedent rectal and genitourinary cultures [19]. They found evidence of an in-frame *aniA* gene in MenC similar to that present in *Neisseria gonorrhea* that confers the ability to survive under anaerobic conditions [19]. Following a large cluster of *N. meningitidis* urethritis in heterosexual men in Ohio, U.S., WGS identified an insertion sequence that had replaced much of the *cps* capsular locus of a MenC strain, rendering it unencapsulated and non-groupable [20].

We sequenced a collection of archived *N. meningitidis* isolates’ genomes from NYC from 2003-2013 and analyzed phylogenetic relationships of the isolates based on core genome alignment, patient risk factors, and demographics to explore spatial connections among cases not evident from the epidemiologic data alone. The study was undertaken to better understand how IMD transmission occurs within a large urban center with the goal of improving IMD outbreak response, prevention, and control.

## MATERIALS AND METHODS

### Study Population and Case Investigation

IMD is a reportable condition in NYC, where all cases are investigated to identify close contacts who qualify for post-exposure prophylaxis and to uncover links that could indicate an outbreak. Medical providers and laboratories are required to report suspected and confirmed cases within 24 hours as defined by the Council of State and Territorial Epidemiologists (CSTE)/Centers for Disease Control and Prevention’s (CDC) case definition [21]. Sterile site isolates are sent to the NYC Public Health Laboratory for serogrouping and molecular typing. A significant proportion of IMD cases have negative cultures [22]. Beginning in 2006, clinical samples were routinely sought on culture-negative cases for RT-PCR testing at the NY State Wadsworth Center [23]. Non-sterile *N. meningitidis* reports (pharyngitis, urethritis, proctitis) were occasionally received and processed as above on a case-by-case basis. Isolates from non-NYC residents who had an epidemiologic link to NYC were sought for molecular characterization and comparison. Ob1-associated cases were defined as those meeting the CSTE/CDC case definition with *N. meningitidis* serogroup C and PFGE findings of ≥ 85% similarity to the designated outbreak strain, or for probable cases (culture negative), patients who were household contacts to or shared an epidemiologic link with an outbreak-associated case [2]. Ob2-associated cases were defined as confirmed or probable *N. meningitidis* serogroup C MSM NYC residents with onset during August 2010-February 2013 [3].

IMD cases were investigated using a structured data collection instrument. Death from IMD was defined as occurring within 30 days of diagnosis and was obtained through a data match with the NYC Office of Vital Statistics death registry; HIV status was determined through matching with the NYC HIV Surveillance Registry. The earliest available MenC isolate was from 2003, therefore, isolates from 2003-2013 were eligible for inclusion in the study. A selection of other serogroups, non-NYC residents, and non-sterile sources were included for comparison.

### Isolate regrowth, DNA extraction, sequencing, assembly, and annotation

*N. meningitidis* isolates were preserved in Trypticase soy broth with 20% glycol at −70°C. Isolates were removed from storage and plated on Chocolate II Agar (BBL). Following overnight growth at 35° in a humidified 5% CO_2_ incubator, single colonies were selected to inoculate a subculture. DNA was extracted from fresh *N. meningitidis* subculture colonies using Genomic-tip 20/G (Qiagen). Post-extraction quality control was performed to ensure concentration, purity, integrity, and sterility. WGS was performed using the HiSeq 2000 platform (Illumina) at the Edgewood Chemical and Biological Center (ECBC) sequencing facility. DNA was prepared for sequencing using the Nextera DNA Library Preparation kit (Illumina) and paired-end sequencing was performed on the dual-indexed isolate libraries (Illumina) according to the manufacturer’s protocol. After sequence capture, reads were assembled *de novo* using either Velvet (1.2.10) [24] for higher coverage read files (>200Mb) or CLC Genomics Workbench (6.5.1) (Qiagen) for lower coverage read files (<200Mb). Consensus genomes were then annotated using RAST toolkit (1.3.0) [25, 26]. Sequencing personnel were blinded to the epidemiological and geographic metadata.

### Determining orthologs, core genome, and pan genome

Ortholog clusters were determined based on the comparison of all protein coding sequences with OrthoMCL (version 2.0.9) [27, 28] with an e-value cut off of 10^-5^ on the BLAST+ (version 2.27) [29]. The output was then parsed with custom Python scripts to find the core genome for different sets of isolates and genes specific to only isolates with particular metadata characteristics. The ortholog analysis also included 10 additional finished *N. meningitidis* genomes from GenBank and one draft genome from Paris reported to be a part of the MSM outbreak [30]. Assembled genomes were queried using BLAST against the PubMLST *Neisseria* database (https://pubmlst.org/neisseria/) to identify *N. meningitidis* clonal complex and the presence of a frameshift mutation in the *aniA* gene [31].

### Phylogeny

The phylogeny of the genomes included in the OrthoMCL analysis was determined based on the core genome. We used a strict definition of core genome as being those ortholog clusters that contain only a single gene from each genome, excluding clusters with any paralogs. No epidemiological data were taken into consideration for determining orthologs or the initial phylogeny. The DNA sequences for each cluster were individually aligned using MAFFT (version 7.130b) [32], then Trimal (version 1.2) [33] was used to obtain the mean percent identity from each individual alignment. Alignments with a mean percent identity less than 90% were inspected individually and those with poor alignments, suggesting that they may not be orthologs, were removed from further analysis. The remaining alignments were concatenated into one core genome alignment with FASconCAT (version 1.02) [34] to be used by RAxML (version 8) [35] to determine phylogenetic relationships using maximum likelihood optimality criterion. The same methods were used for determining the core genomes of a subset of genomes for analyses (i.e., Genome Wide Association Studies and Betweenness centrality (see below)).

### Genome Wide Association Studies

Genome Wide Association Studies (GWAS) were done on two different input data types: (1) gene present or absent in a genome; (2) single nucleotide polymorphisms (SNPs) in the core genome. Custom Python scripts were written to parse the OrthoMCL output to determine if a gene was present or absent in each genome based on whether there was a gene listed within orthologous clusters and to produce PED and MAP files. Therefore, the number of genes considered is equivalent to the total number of gene clusters determined by OrthoMCL, 5,280 gene clusters. In the PED file, one base was assigned if the gene was present and a different base used if the gene was absent for that genome for each orthologous cluster. Custom python scripts were written to find SNPs in alignments made by MAFFT for each individual gene cluster of the strict core genome and to produce PED and MAP files. Those alignment locations that contained more than two SNP variations were excluded due to limitations in the GWAS software, but the presence of an N was included as missing data for that genome. The program PLINK (version 1.07)[36] was then used to perform the GWAS with the genomic location set to the haploid mitochondria in the input files. P-values were adjusted by PLINK for population stratification using a genomic control and for multiple testing using a conservative Bonferroni correction. Gene Ontology (GO) terms were determined by BLAST2GO [37].

### Betweenness centrality

For the betweenness centrality analysis we used methods similar to those used in Janies et al. [38]. We used the subset of isolates with home addresses that could be geocoded plus an isolate from Paris, LNP27256 (GenBank: AVOU01000001.1) related to the outbreak [31]. When multiple isolates originated from the same patient, one isolate was selected as a representative. We then used the phylogenetic methods describe above to produce a tree based on the strict core genome for only these isolates [39]. When RAxML was run on the concatenated core genome, the earliest isolate, S00-1196, was used as an outgroup. This tree was then brought into PAUP* (version 4.0a147) with the cases' home borough or neighborhood as units of a geography. Borough was determined from the latitude and longitude of the home address using ArcGIS (http://www.esri.com/software/arcgis/explorer/index.html). New York City neighborhood was determined based on NYC Department of City Planning Neighborhood Tabulation Areas (NTA, see Table S1). Isolates from outside NYC maintained their individual location for both analyses. PAUP* was then used to calculate the ancestor descendent changes in geographic location at each branch of the phylogenetic tree. The directionality and number of each type of change was then measured. These data were then integrated to form the network of bacterial transmission using igraph and qgraph packages in R [40, 41].

### Ethical Statement

The Department of Health and Mental Hygiene (DOHMH) Institutional Review Board determined the study to be public health surveillance that is non-research.

## RESULTS

### Study Population and Isolate Sequencing

A total of 102 isolates were submitted for WGS. Six isolates were excluded due to poor sequencing or assembly of their genomes and there were 12 repeat isolates (same patient, different specimen source) from 11 individuals leaving 84 unique patients (74 NYC residents, 63 invasive infections) in the analysis (see Figure 1). During the years 2003-2013, a total of 322 IMD cases were reported in NYC residents, of which 263 were culture positive (the universe of possible study isolates). The proportion of eligible invasive isolates in NYC residents included in the study by serogroup is as follows: 52/86 (60%) MenC, 4/60 (7%) MenB, 4/21 (19%) MenW, and 3/89 (3%) MenY. Fourteen MenC patient isolates were included from each outbreak (Ob1 had 21 and Ob2 had 15 culture positive cases). The demographic characteristics of invasive MenC isolates from NYC residents included in the study were compared to MenC patients during the same interval whose isolates were unavailable (includes culture negative cases that met the case definition). Patients with MenC included in the study were more likely to be older, male and non-Hispanic (Table 1). Individual patient and isolate characteristics are presented in Table 2.

**Figure. 1.**
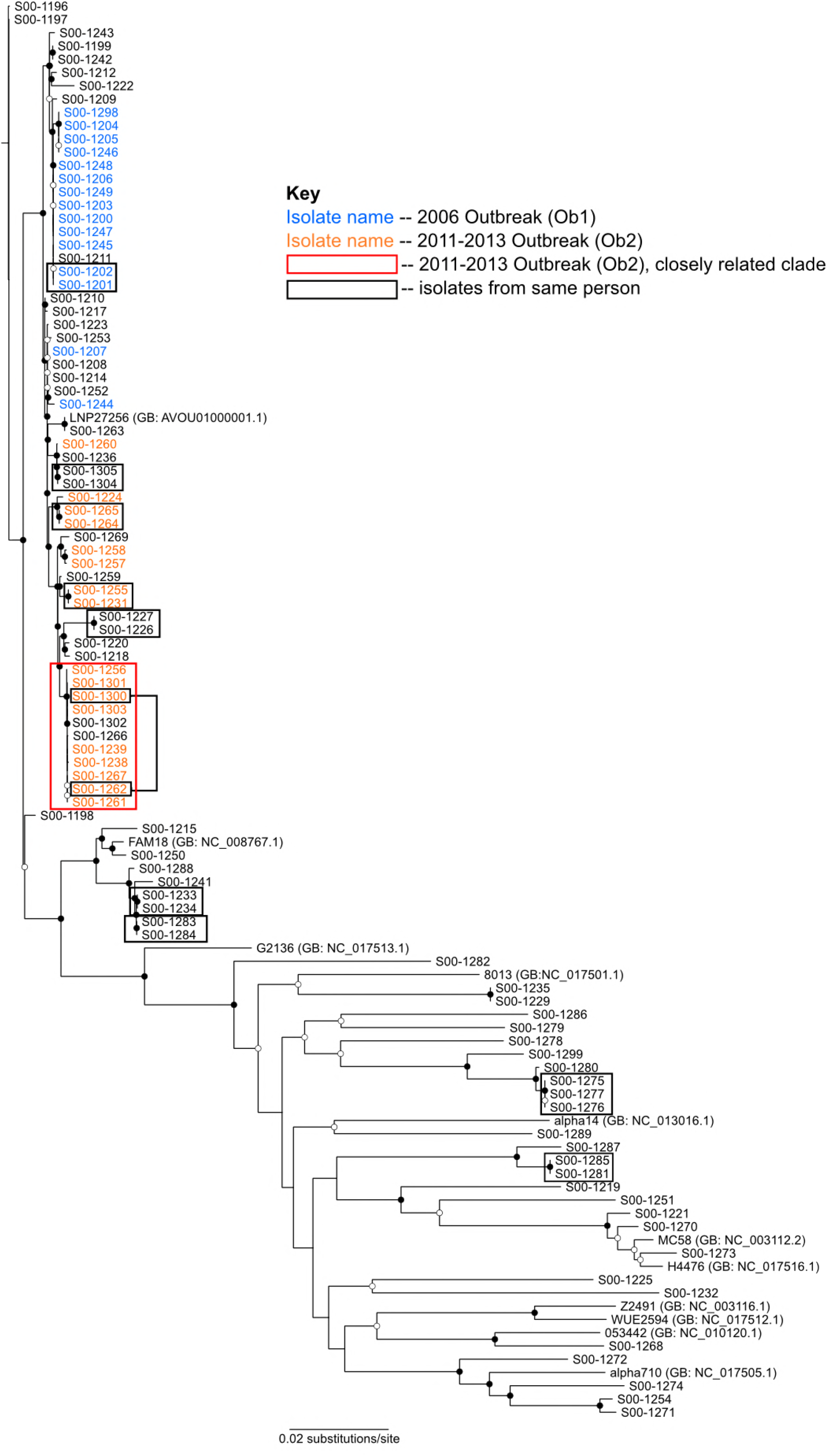
Phylogenetic relationship of *N. meningitidis* isolates from around NYC 2003-2013. The tree is based on core genome SNPs and rooted on the oldest isolate. Open circles represent nodes with bootstrap support ≥50, while closed circles represent nodes with bootstrap support ≥95 based on 100 replicates. Those isolates that are part of the 2006 (Ob1) based on epidemiology outbreak have blue text. Those isolates that are part of the 2012 outbreak (Ob2) based on epidemiology have orange text, with the closely related subclade indicated by a red box. Isolates from the same individuals are indicated by black boxes.

**Table 1.** Table 1-Characteristics of Sequenced Invasive MenC isolates compared to those that were unavailable (confirmed and probable IMD cases), New York City, 2003–2013.

| Characteristic | Not Sequenced (n=49) | Sequenced (n=52) |
| --- | --- | --- |
| Age group |  |  |
| 0–4 y | 3 (6.1) | 2 (3.9) |
| 5–14 y | 3 (6.1) | 5 (9.6) |
| 15–24 y | 11 (22.5) | 4 (7.7) |
| 25–44 y | 20 (40.8) | 20 (38.5) |
| 45–64 y | 7 (14.3) | 14 (26.9) |
| ≥65 y | 5 (10.2) | 7 (13.5) |
| Sex |  |  |
| Female | 23 (46.9) | 19 (36.5) |
| Male | 26 (53.1) | 33 (63.5) |
| Race/ethnicity |  |  |
| Non-Hispanic white | 7 (14.3) | 14 (26.9) |
| Non-Hispanic black | 16 (32.7) | 23 (44.2) |
| Hispanic | 16 (32.7) | 9 (17.3) |
| Asian/Pacific Islander | 2 (4.1) | 2 (3.9) |
| Unknown/Other | 8 (16.3) | 4 (7.7) |
| Borough of residence |  |  |
| Bronx | 7 (14.3) | 7 (13.5) |
| Brooklyn | 27 (55.1) | 26 (50.0) |
| Manhattan | 9 (18.3) | 13 (25.0) |
| Queens | 5 (10.2) | 5 (9.6) |
| Staten Island | 1 (2.0) | 1 (1.9) |
| HIV | 12 (24.5) | 10 (19.2) |
| MSM (Males, 2011–2013, age≥15) | 8/9 (88.9) | 13/15 (86.7) |
| Died | 12 (24.5) | 22 (42.3) |
| Outbreak 1 (2005-2006) | 9/23 (39.1) | 14/23 (60.9) |
| Outbreak 2 (2010-2013) | 7/22 (31.8) | 14/22 (63.6) |

**Table 2.** Individual Patient and Isolate Characteristics of *Neisseria meningitidis***Strains Included in the Study**.

| Patient ID | Study ID | Month/Year | Age (Years) | Sex | Out-break | S G | Residency | Specimen Source | Sequence type/ Clonal Complex | In-Frame *aniA* |
| --- | --- | --- | --- | --- | --- | --- | --- | --- | --- | --- |
| 1 | S00-1196 | Jan2003 | 25 | M | 0 | C | Brooklyn | Blood | ST-11 complex/ET-37 complex | No |
| 2 | S00-1197 | Nov2003 | 32 | M | 0 | C | Suffolk County, NY | Blood | ST-11 complex/ET-37 complex | No |
| 3 | S00-1198 | Jan2004 | 46 | F | 0 | C | Brooklyn | Blood | ST-11 complex/ET-37 complex | Yes |
| 4 | S00-1199 | Jun2004 | 49 | F | 0 | C | Manhattan | Blood | ST-11 complex/ET-37 complex | Yes |
| 5 | S00-1200 | Nov2005 | 49 | F | 1 | C | Bronx | CSF | ST-11 complex/ET-37 complex | No |
| 6 | S00-1201 | Jan2006 | 40 | M | 1 | C | Brooklyn | Blood | ST-11 complex/ET-37 complex | No |
| 6 | S00-1202 | Jan2006 | 40 | M | 1 | C | Brooklyn | CSF | ST-11 complex/ET-37 complex | No |
| 7 | S00-1203 | Feb2006 | 66 | M | 1 | C | Brooklyn | Blood | ST-11 complex/ET-37 complex | No |
| 8 | S00-1204 | Mar2006 | 49 | M | 1 | C | Brooklyn | Blood | ST-11 complex/ET-37 complex | No |
| 9 | S00-1205 | Apr2006 | 41 | M | 1 | C | Brooklyn | Blood | ST-11 complex/ET-37 complex | No |
| 10 | S00-1206 | Aug2006 | 43 | M | 1 | C | Brooklyn | Blood | ST-11 complex/ET-37 complex | No |
| 11 | S00-1207 | Nov2006 | 30 | F | 1 | C | Brooklyn | Rectal | ST-11 complex/ET-37 complex | Yes |
| 12 | S00-1208 | Jan2007 | 47 | M | 0 | C | Manhattan | Rectal | ST-11 complex/ET-37 complex | Yes |
| 13 | S00-1209 | May2007 | 41 | M | 0 | C | Suffolk County, NY | Pericardial | ST-11 complex/ET-37 complex | Yes |
| 14 | S00-1210 | Jun2007 | 73 | M | 0 | C | Brooklyn | Blood | ST-11 complex/ET-37 complex | Yes |
| 15 | S00-1211 | Jun2007 | 61 | F | 0 | C | Manhattan | Blood | ST-11 complex/ET-37 complex | No |
| 16 | S00-1212 | Aug2007 | 0 | F | 0 | C | Staten Island | Blood | ST-11 complex/ET-37 complex | No |
| 17 | S00-1214 | Dec2007 | 65 | M | 0 | C | Manhattan | Blood | ST-11 complex/ET-37 complex | Yes |
| 18 | S00-1215 | Mar2008 | 81 | M | 0 | C | Manhattan | CSF | ST-11 complex/ET-37 complex | No |
| 19 | S00-1217 | Jun2008 | 58 | M | 0 | C | Manhattan | Blood | ST-11 complex/ET-37 complex | Yes |
| 20 | S00-1218 | Aug2008 | 36 | M | 0 | C | Bronx | Blood | ST-11 complex/ET-37 complex | Yes |
| 21 | S00-1219 | Sep2008 | 83 | F | 0 | C | Queens | Blood | ST-35 complex | Yes |
| 22 | S00-1220 | Sep2008 | 58 | F | 0 | C | Brooklyn | CSF | ST-11 complex/ET-37 complex | Yes |
| 23 | S00-1221 | Apr2010 | 5 | M | 0 | C | Queens | Blood | ST-32 complex/ET-5 complex | Yes |
| 24 | S00-1222 | May2010 | 85 | F | 0 | C | Brooklyn | Blood | ST-11 complex/ET-37 complex | No |
| 25 | S00-1223 | Jul2010 | 1 | M | 0 | C | Chesterfield County, South Carolina | Blood | ST-11 complex/ET-37 complex | Yes |
| 26 | S00-1224 | Aug2010 | 47 | M | 2 | C | Manhattan | CSF | ST-11 complex/ET-37 complex | Yes |
| 27 | S00-1225 | Oct2010 | 20 | M | 0 | C | Manhattan | Rectal | Not defined | Yes |
| 28 | S00-1226 | Nov2010 | 44 | F | 0 | C | London, England | CSF | ST-11 complex/ET-37 complex | Yes |
| 28 | S00-1227 | Nov2010 | 44 | F | 0 | C | London, England | Blood | ST-11 complex/ET-37 complex | Yes |
| 29 | S00-1229 | Jan2011 | 4 | M | 0 | C | Bronx | Blood | ST-103 complex | Yes |
| 30 | S00-1231 | Aug2011 | 31 | M | 2 | C | Brooklyn | Blood | ST-11 complex/ET-37 complex | Yes |
| 30 | S00-1255 | Aug2011 | 31 | M | 2 | C | Brooklyn | CSF | ST-11 complex/ET-37 complex | Yes |
| 31 | S00-1232 | Sep2011 | 17 | M | 0 | C | Suffolk County, NY | Trachea | Not defined | Yes |
| 32 | S00-1233 | Oct2011 | 22 | M | 0 | C | Brooklyn | Rectal | ST-11 complex/ET-37 complex | No |
| 32 | S00-1234 | Oct2011 | 22 | M | 0 | C | Brooklyn | Rectal | ST-11 complex/ET-37 complex | No |
| 33 | S00-1235 | Dec2011 | 35 | M | 0 | C | Brooklyn | Blood | ST-103 complex | Yes |
| 34 | S00-1236 | Mar2012 | 49 | M | 0 | C | Queens | Blood | ST-11 complex/ET-37 complex | Yes |
| 35 | S00-1238 | Nov2012 | 33 | M | 2 | C | Brooklyn | CSF | ST-11 complex/ET-37 complex | Yes |
| 36 | S00-1239 | Feb2013 | 53 | M | 2 | C | Manhattan | CSF | ST-11 complex/ET-37 complex | Yes |
| 37 | S00-1241 | Apr2013 | 57 | F | 0 | W | Bronx | Blood | ST-11 complex/ET-37 complex | No |
| 38 | S00-1242 | Dec2003 | 22 | F | 0 | C | Bronx | Blood | ST-11 complex/ET-37 complex | Yes |
| 39 | S00-1243 | Jan2006 | 25 | M | 0 | C | Manhattan | Rectal | ST-11 complex/ET-37 complex | No |
| 40 | S00-1244 | Jan2006 | 6 | M | 1 | C | Brooklyn | Blood | ST-11 complex/ET-37 complex | Yes |
| 41 | S00-1245 | Feb2006 | 43 | F | 1 | C | Brooklyn | Blood | ST-11 complex/ET-37 complex | No |
| 42 | S00-1246 | Apr2006 | 13 | M | 1 | C | Brooklyn | CSF | ST-11 complex/ET-37 complex | No |
| 43 | S00-1247 | Apr2006 | 43 | F | 1 | C | Brooklyn | Blood | ST-11 complex/ET-37 complex | No |
| 44 | S00-1248 | Jun2006 | 46 | F | 1 | C | Brooklyn | Blood | ST-11 complex/ET-37 complex | No |
| 45 | S00-1249 | Jun2006 | 12 | F | 1 | C | Brooklyn | Blood | ST-11 complex/ET-37 complex | No |
| 46 | S00-1250 | Jul2007 | 44 | M | 0 | C | Manhattan | Blood | ST-11 complex/ET-37 complex | No |
| 47 | S00-1251 | Jan2008 | 30 | M | 0 | C | Manhattan | Rectal | ST-269 complex | No |
| 48 | S00-1252 | May2008 | 42 | F | 0 | C | Manhattan | Blood | ST-11 complex/ET-37 complex | Yes |
| 49 | S00-1253 | Jul2008 | 20 | F | 0 | C | Bronx | Blood | N/A | Yes |
| 50 | S00-1254 | Jul2009 | 19 | M | 0 | N<br>O<br>N | Bronx | Rectal | ST-41/44 complex/Lineage 3 | Yes |
| 51 | S00-1256 | Aug2011 | 52 | M | 2 | C | Manhattan | Blood | ST-11 complex/ET-37 complex | Yes |
| 52 | S00-1257 | Jan2012 | 23 | M | 2 | C | Brooklyn | Blood | ST-11 complex/ET-37 complex | No |
| 53 | S00-1258 | Mar2012 | 28 | M | 2 | C | Brooklyn | Blood | ST-11 complex/ET-37 complex | No |
| 54 | S00-1259 | Mar2012 | 65 | F | 0 | C | Manhattan | Blood | ST-11 complex/ET-37 complex | Yes |
| 55 | S00-1260 | Jun2012 | 59 | M | 2 | C | Brooklyn | Blood | ST-11 complex/ET-37 complex | Yes |
| 56 | S00-1261 | Sep2012 | 42 | M | 2 | C | Bronx | Blood | ST-11 complex/ET-37 complex | Yes |
| 57 | S00-1262 | Sep2012 | 32 | M | 2 | C | Manhattan | CSF | ST-11 complex/ET-37 complex | Yes |
| 57 | S00-1300 | Sep2012 | 32 | M | 2 | C | Manhattan | CSF | ST-11 complex/ET-37 complex | Yes |
| 58 | S00-1263 | Nov2012 | 20 | M | 0 | C | Philadelphia, PA | Blood | ST-11 complex/ET-37 complex | Yes |
| 59 | S00-1264 | Dec2012 | 33 | M | 2 | C | Manhattan | Blood | ST-11 complex/ET-37 complex | Yes |
| 59 | S00-1265 | Dec2012 | 33 | M | 2 | C | Manhattan | Blood | ST-11 complex/ET-37 complex | Yes |
| 60 | S00-1266 | Dec2012 | 29 | M | 0 | C | Middlesex County, NJ | Blood | ST-11 complex/ET-37 complex | Yes |
| 61 | S00-1267 | Feb2013 | 26 | M | 2 | C | Brooklyn | Blood | ST-11 complex/ET-37 complex | Yes |
| 62 | S00-1268 | Apr2013 | 41 | M | 0 | B | Manhattan | Throat | ST-4821 complex | Yes |
| 63 | S00-1269 | Apr2013 | 31 | M | 0 | C | Essex County, NJ | CSF | ST-11 complex/ET-37 complex | No |
| 64 | S00-1270 | Dec2011 | 40 | M | 0 | B | Bronx | Blood | ST-32 complex/ET-5 complex | Yes |
| 65 | S00-1271 | Dec2011 | 0 | M | 0 | B | Queens | Blood | ST-41/44 complex/Lineage 3 | Yes |
| 65 | S00-1273 | Apr2012 | 24 | F | 0 | B | Manhattan | CSF | ST-32 complex/ET-5 complex | Yes |
| 66 | S00-1272 | Aug2011 | 0 | M | 0 | B | Brooklyn | Blood | N/A | Yes |
| 67 | S00-1274 | Jan2013 | 6 | F | 0 | B | Westchester County, NY | Blood | N/A | Yes |
| 68 | S00-1275 | Jan2010 | 0 | M | 0 | Y | Bronx | CSF | ST-23 complex/Cluster A3 | No |
| 68 | S00-1276 | Jan2010 | 0 | M | 0 | Y | Bronx | Blood | ST-23 complex/Cluster A3 | No |
| 68 | S00-1277 | Jan2010 | 0 | M | 0 | Y | Bronx | Blood | ST-23 complex/Cluster A3 | No |
| 69 | S00-1278 | Jul2008 | 86 | F | 0 | Y | Brooklyn | Blood | ST-174 complex | Yes |
| 70 | S00-1279 | Nov2012 | 24 | M | 0 | Y | Bronx | Blood | ST-167 complex | No |
| 71 | S00-1280 | May2013 | 48 | M | 0 | Y | Bronx | Trachea | ST-23 complex/Cluster A3 | No |
| 72 | S00-1281 | Apr2010 | 22 | M | 0 | W | Manhattan | Blood | ST-1157 complex | Yes |
| 72 | S00-1285 | Apr2010 | 22 | M | 0 | W | Manhattan | CSF | ST-1157 complex | Yes |
| 73 | S00-1282 | Jun2012 | 53 | F | 0 | W | Bronx | Blood | ST-22 complex | Yes |
| 74 | S00-1283 | Apr2009 | 27 | M | 0 | W | Queens | Blood | ST-11 complex/ET-37 complex | No |
| 74 | S00-1284 | Apr2009 | 27 | M | 0 | W | Queens | Blood | ST-11 complex/ET-37 complex | No |
| 75 | S00-1286 | Nov2012 | 24 | M | 0 | W | Manhattan | Throat | N/A | Yes |
| 76 | S00-1287 | Apr2013 | 38 | M | 0 | N O N | Manhattan | Throat | ST-1157 complex | Yes |
| 77 | S00-1288 | May2013 | 27 | M | 0 | W | Manhattan | Throat | ST-11 complex/ET-37 complex | No |
| 78 | S00-1289 | May2013 | 68 | M | 0 | B | Brooklyn | Blood | ST-213 complex | Yes |
| 79 | S00-1298 | Oct2006 | 6 | M | 1 | C | Brooklyn | Blood | ST-11 complex/ET-37 complex | No |
| 80 | S00-1299 | Aug2008 | 60 | F | 0 | C | Bronx | Blood | ST-23 complex/Cluster A3 | No |
| 81 | S00-1301 | Oct2012 | 27 | M | 2 | C | Queens | Blood | ST-11 complex/ET-37 complex | Yes |
| 82 | S00-1302 | Oct2012 | 47 | M | 0 | C | Suffolk County, NY | Blood | ST-11 complex/ET-37 complex | Yes |
| 83 | S00-1303 | Dec2012 | 23 | M | 2 | C | Brooklyn | Blood | ST-11 complex/ET-37 complex | Yes |
| 84 | S00-1304 | Jun2013 | 43 | F | 0 | C | Queens | Blood | ST-11 complex/ET-37 complex | Yes |
| 84 | S00-1305 | Jun2013 | 43 | F | 0 | C | Queens | Blood | ST-11<br>complex/ET-37 complex | Yes |

Outbreak 1=1, Outbreak 2=2, not part of an outbreak=0; SG=serogroup, N/A= MLST sequence type and clonal complex indeterminate

Our assembled draft genomes had between 218 and 1223 contigs with a mean contig number of 494 and read depth coverage ranging from 300 to 3000 (Table S2). The total genome length and gene count of these contigs are all within the expected size for *N. meningitidis.* The strict core genome, which explicitly left out orthologous clusters containing paralogs, was determined for the 96 genomes sequenced and 11 genomes from GenBank was found to contain 1,171 genes. When only the genomes sequenced in this study were considered for the core genome, it contains 1,373 genes. Since these were draft genomes, it was suggested that a better metric for determining the core genome would be to include gene clusters found in 95% of the genomes being analyzed [42]. Using this metric, the size of the core genome increased to 1,348 genes for 107 genomes and 1,545 genes for only the genomes sequenced in this study. The number of unique genes ranged from 7 to 376 (mean 33) per isolate (Table 2S). The estimated *N. meningitidis* pan genome based on these isolates contained 8,453 genes.

### Core Genome Phylogeny

The phylogenetic relationships among the isolates was determined with maximum likelihood tree searches based on alignments of the strict core genome. From this analysis we determined that Ob1 isolates classified by epidemiology were mostly found on a single clade, with S00-1207 and S00-1244 as the exceptions (Figure 1, blue text). Likewise, the 14 Ob2 isolates mostly formed a separate, single clade, with only S00-1260 found on a different clade (Figure 1, orange text). One Ob2 subclade with dates ranging from August 2011 to February 2013 showed less diversity (Figure 1, 11 isolates, outlined in red) and included two patients who were not NYC residents (S00-1266 and S00-1302). All of the patients in the subclade were male between the ages of 23 and 53 years; and of the 9 NYC residents: seven were HIV-infected, three died, and seven reported drug use (methamphetamine, cocaine, and/or marijuana). The subclade contained 150 SNPs within the core genome, compared to 4,921 core genome SNPs in the entire outbreak, indicating that the isolates in the subclade were more closely related to one another than to the rest of the outbreak isolates. Of the eight isolates found within Ob2’s clade that were not declared to be part of this outbreak based on epidemiology, six (S00-1269, S00-1259, S00-1227, S00-1226, S00-1302, and S00-1266) were collected during outbreak’s time period, with S00- 1220 and S00-1218 as the exceptions collected in 2008, before Ob2 began (Table 2 and Figure 1). Non-resident patients S00-1266, S00-1269, and S00-1302 were not known to be MSM and were not considered part of Ob2 at the time of collection. Due to the isolates’ position in the tree, they may have shared a common exposure with outbreak cases or their social network. Isolate S00-1259 was obtained from a 65 year-old woman in March 2012. She reported no epidemiologic link to the outbreak and was not included as an Ob2 case. While the isolate’s position on the tree suggests inclusion as part of Ob2, it is also possible that the case represents sporadic transmission.

### Presence of In-frame *aniA* Gene

Fifty-seven percent (55/96) of isolates and 57% of patients (48/84) contained an in-frame *aniA* gene coding for nitrite reductase. All multiple isolates were concordant. The earliest appearance of an in-frame *aniA* was in 2003 (Table 2) and the proportion by unique patient increased over the interval from 45% (17/38) in the years 2003-2008 to 67% (31/46) in the years 2009-2013. The proportion of isolates with an intact *aniA* gene increased from 14% (2/14) patients for Ob1 to 86% (12/14) for Ob2. All 11 Ob2 subclade isolates contained the in-frame *aniA* gene.

### Genome Wide Association Studies

Genome Wide Association Studies were performed using only the genomes sequenced for the study using two metrics: (1) the presence or absence of genes; and (2) the presence of SNPs in the strict core genome. Due to software limitations, only SNPs with 2 variants could be considered. We examined 82,597 SNPs, while having to exclude 3,541 SNP locations from the analysis. We tested for the association of SNPs or genes to both Ob1 and Ob2 (Table 3). The analysis for associations with Ob2 excluded S00-1260 due to its phylogenetic distance from the rest of the Ob2 isolates.

**Table 3.**
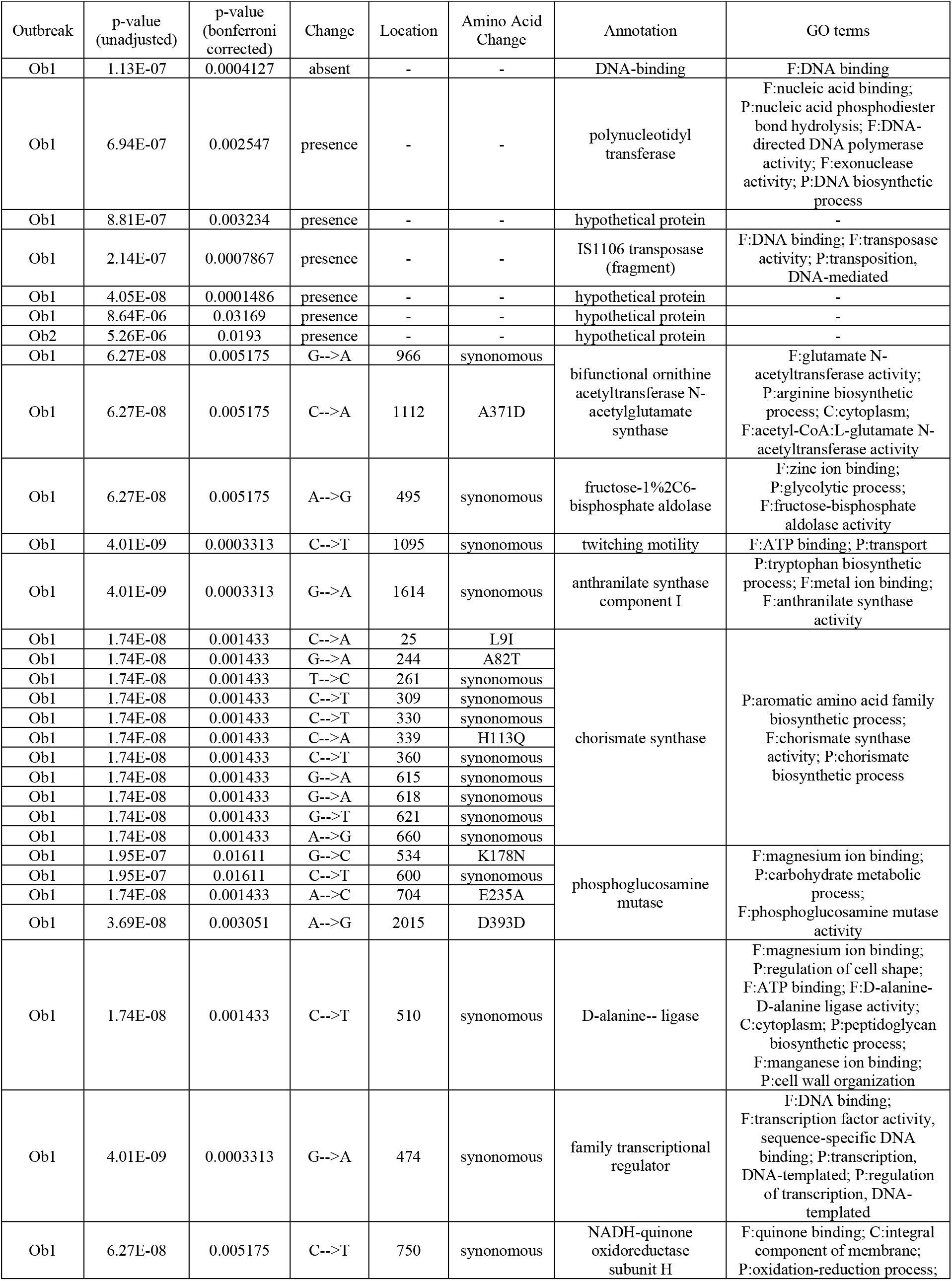
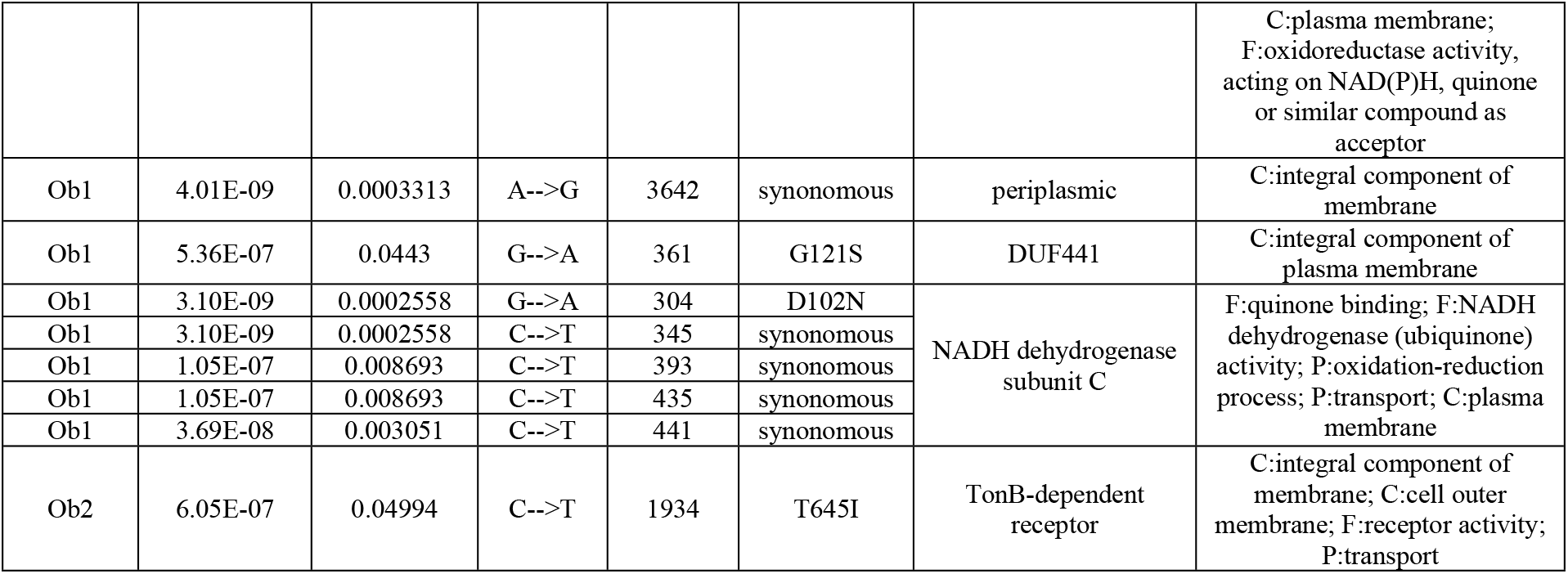
Genome Wide Association Studies of 96 Sequenced *N. meningitidis* Isolates.

We found the presence of five genes and the absence of one to be significantly associated with Ob1 (p<0.05, Table 3). Ob1 also had 30 SNPs significantly associated with it (p<0.05, Table 3). GO terms indicated that these SNPs are found in genes of a wide variety functions.

Ob2 isolates have one gene encoding a protein associated with the outbreak clade which was annotated as a hypothetical protein (p=0.0193, Table 3). One SNP in a TonB-dependent receptor was found to be significantly associated with Ob2 (p=0.04994, Table 3). This SNP is non-synonymous and results in a change from threonine to isoleucine. All eight isolates that were not epidemiologically considered part of Ob2, but were present on the same clade also possess this change. Through a BLAST search, we found that this SNP is not unique to this clade of *N. meningitidis*.

### Betweenness Centrality

As detailed in methods, we optimized the home location of the patients on the phylogeny and converted the result of all the ancestor descendent changes to a network to determine how *N. meningitidis* moved around the city over the 11-year period. We then applied betweenness centrality as of a measure of how often a node in a network is on the shortest path between two other nodes. Seventy-eight unique patient isolates, with a strict core genome of 1320 genes, were included in the betweenness centrality analysis. At the NYC borough resolution, not all of the boroughs were connected to each other nor are all the connections bidirectional (Figure 2). Brooklyn had the highest betweenness centrality as the only borough connected to all other boroughs and the one with the most connections to non-NYC locations. Brooklyn also had the most intra-borough transmission events (Figure 2, indicated by the arrow that loops back). The Bronx had the second highest betweenness centrality of the boroughs. The Bronx was connected to all the boroughs except Staten Island. We observed bidirectional movement between the Bronx, Brooklyn and Queens, and connections to three non-NYC locations.

**Figure. 2.**
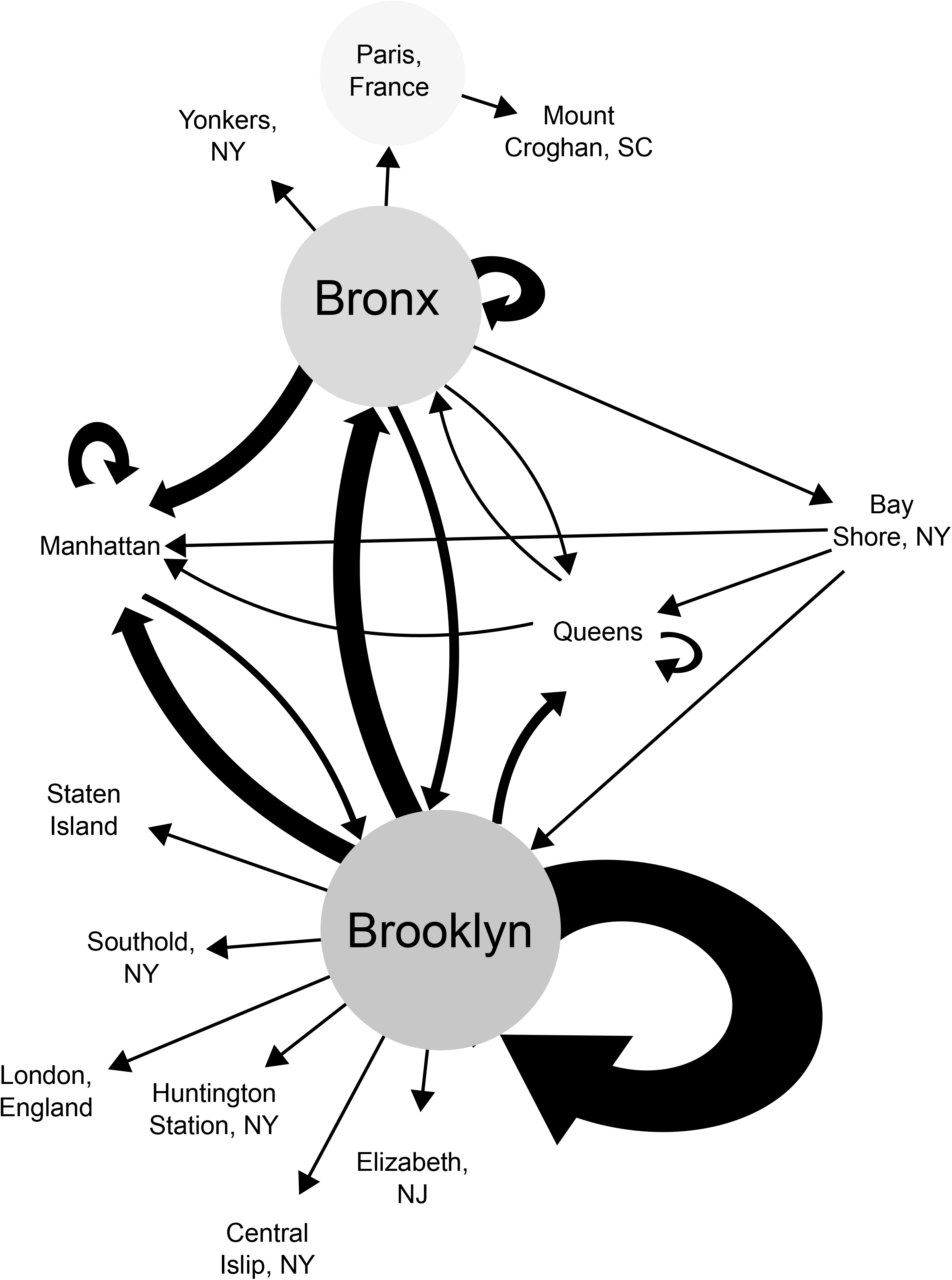
Betweenness and the Movement of *N. meningitidis* around NYC Boroughs. Locations with higher betweenness values have a darker colored circle. The arrows indicate the direction of movement with the size of the arrowheads and thickness of arrow scaling with the amount of movement between those locations, with a larger size indicating greater movement.

While betweenness centrality at the level of the borough allowed us to see an additive summary of *N. meningitidis* movement over 11 years, it did not elucidate the details of transmission events for the two outbreaks. When the betweenness centrality analysis was examined at the level of neighborhood, the arrows represented much more granular transmission events between locations elucidating the transmissions specific to each outbreak (Figure 3). Most transmission events at the level of neighborhood were single events. However, there were some multiple transmissions between or within locations: (1) two transmissions occurred between Stuyvesant Heights (BK35) and Bedford (BK75); (2) three transmissions occurred within Stuyvesant Heights (BK35); and (3) two transmissions occurred within Crown Heights North (BK61) (Figure 3). All three of these multiple transmission events were from Ob1 and only two non-Brooklyn neighborhoods were involved with spread which establishes the geographic concentrations of this outbreak in Brooklyn (Figure 3, colored blue). The two non-Brooklyn neighborhoods involved in Ob1 were the Bronx neighborhood of Co-Op City (BX13) and the Manhattan neighborhood of Carnegie Hill (MN40). The transmission to Co-Op City (BX13) was a dead end for further transmission (Figure 3, colored blue). From Carnegie Hill (MN40), two transmissions occurred to two different Brooklyn neighborhoods that were disconnected from the rest of the Ob1 outbreak (Figure 3, colored blue).

**Figure. 3.**
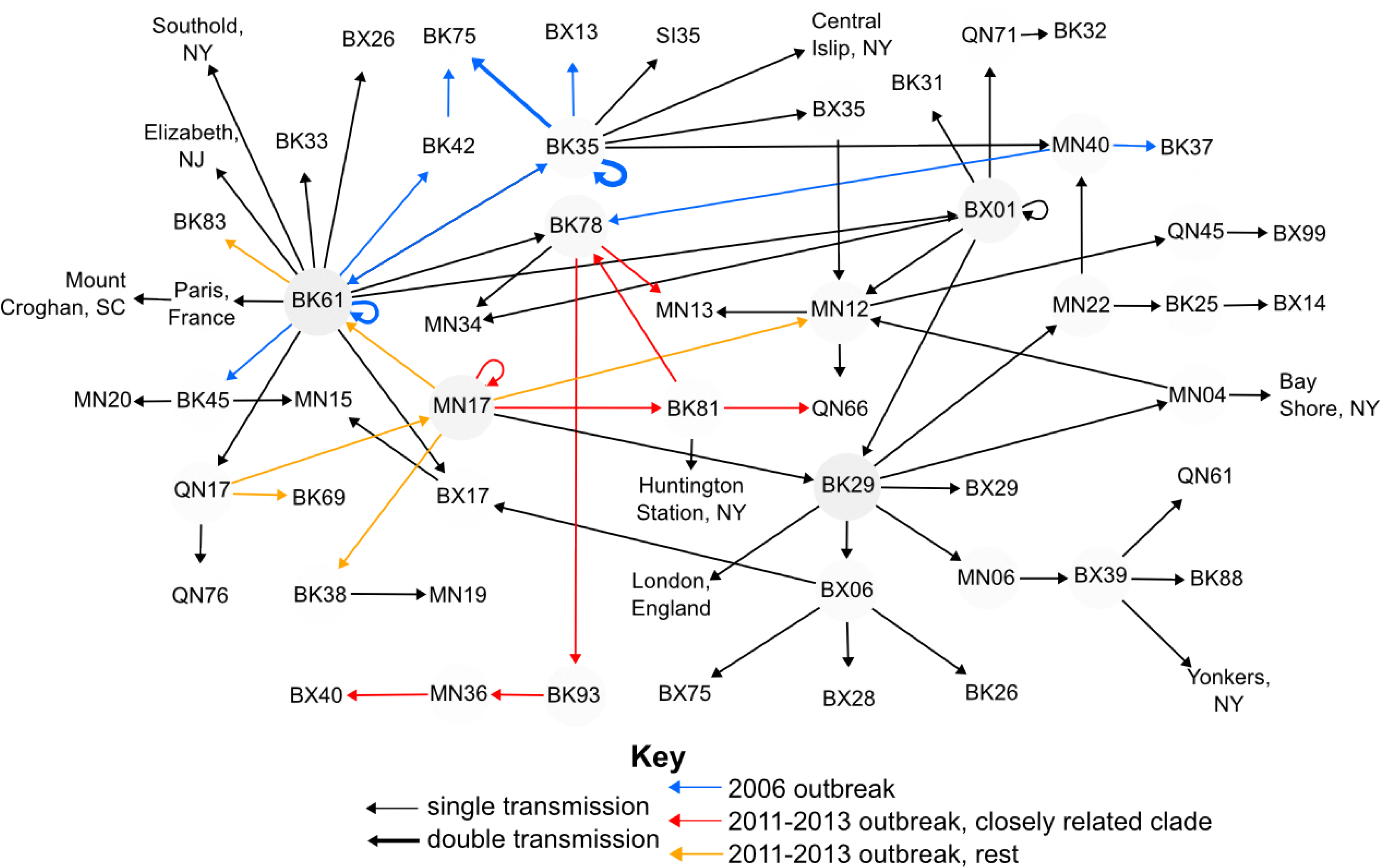
Betweenness and the Movement of *N. meningitidis* around NYC Neighborhoods. Locationswith higher betweenness values have a darker colored circle. Most arrows represents single transmission event between neighborhoods with the 3 exceptions that have slightly thicker arrows: (1) two transmissions happened between BK35 (Stuyvesant Heights) and BK75 (Bedford), (2) two transmissions happened within BK61 (Crown Heights North), (3) three transmissions happened within BK35 (Stuyvesant Heights). Arrows colored blue represent those transmissions that were part Ob1. Ob2 is designated with arrows colored either red for the closely related subclade and orange for the rest of the outbreak isolates. The neighborhood codes are all the official designation from New York City Neighborhood Tabulation Areas. A key for those referenced is in Supplemental Table 1. Those from the same borough start with the same letters; BX:Bronx, BK:Brooklyn, MN:Manhattan, SI:Staten Island, QN:Queens.

For Ob2 Midtown/Midtown South (MN17) is the most important neighborhood for transmission (Figure 3, colored red and orange). The closely related Ob2 clade originates in this neighborhood before moving to Brownsville (BK81) from which the transmission fans out to multiple boroughs.

## DISCUSSION

We identified 102 *N. meningitidis* isolates from 2003-2013 and successfully performed WGS on 96 of them to uncover connections among patients not found during case investigations. Our *de novo* genome assembly metrics were comparable to that of other researchers using similar sequencing methods. Multiple studies have estimated that the core genome of *N. meningitidis* is around 1300-1600 genes [43, 44, 45], which is consistent with what we found for the genomes analyzed here. As expected, the pan genome for this group is larger than what has been seen in previous studies. This is because *N. meningitidis* has an open pan genome with an estimated 38 new genes added to its pan genome with each newly sequenced isolate [43]. Consistent with this estimate, our genomes had an average of 33 unique genes.

The core genome phylogenetic analysis provided evidence that the epidemiologically determined Ob1 and Ob2 were separate outbreaks caused by different, yet related, strains. The overall shape of the phylogenetic tree indicates that while many of the isolates in NYC have highly similar core genomes, including those from Ob1 and Ob2, there are a number of isolates that are very different, having greater similarity to isolates sequenced as parts of other studies. This could be due to the introduction of *N. meningitidis* strains from other parts of the world through immigration and tourism.

Included in the Ob2 clade are two isolates (S00-1218 and S00-1220) from 2008 indicating that isolates with highly similar core genomes were present in NYC prior to the start of this outbreak. These isolates could be missing important genetic elements outside of the core genomes that conferred the Ob2 strain added virulence or afforded it a competitive advantage. Alternatively, and more likely, patient behaviors and social network dynamics differed for those isolates, preventing them from triggering an outbreak in 2008.

Our analysis also identified isolates that should have been considered part of an outbreak based on genetic relatedness that were not epidemiologically categorized as such. For example, S00- 1211 is located with the Ob1 isolates on the phylogenetic tree, however, the patient had onset more than six months after the end of the outbreak, was not known to use drugs, and did not live in Brooklyn. The patient resided in a shelter and her onset was close in time to two other IMD cases associated with shelters and substance abuse (neither isolate was available for WGS), suggesting that it is likely the patient came in contact with a person carrying the Ob1 lineage strain.

During Ob2, one case of serogroup C IMD, indistinguishable by PFGE from Ob2 isolates, occurred in a woman (S00-1259) who denied contact with the MSM community and was therefore not considered part of the outbreak. The position of S00-1259 on the phylogenetic tree suggests that she was infected by the Ob2 strain. Two non-resident male cases (S00-1266 and S00-1302) were also not considered part of the outbreak. One patient died and was not known to family members to be MSM and it was later learned that the other individual reported MSM in the year prior to his IMD. Both of these patient isolates lie within the Ob2 subclade and therefore were likely part of the MSM outbreak. One other MSM outbreak-associated patient isolate (S00- 1260), which was not related by PFGE criteria, appears in a different part of the phylogenetic tree than the Ob2 strains and we now consider this to have been a sporadic case and unrelated to Ob2.

We were able to document the appearance of an intact *aniA* gene in a NYC IMD patient from 2003, and that the proportion of patient isolates with this mutation increased from Ob1 to Ob2. The public health implications of this adaptation are not fully understood, however, two recent observations are of concern: 1) the emergence in the US Midwest of *N. meningitidis* urethritis in heterosexual men; and 2) a NYC newborn with *N. meningitidis* conjunctivitis suspected to have been acquired by vertical transmission. The *N. meningitidis* strain in these patients contained an intact *aniA* gene [20, Kretz et al., submitted].

We were limited to the core genome for the SNP GWAS analysis and therefore were unable to look for SNPs in genes that were present in some, but not all, genomes examined. This may have restricted the identification of SNPs important to the epidemiology of the outbreak. Due to bacterial clonal reproduction, GWAS analyses are complicated by phylogenetic relationships and linkage disequilibrium. For this reason, we included in our analysis correction for population stratification using a genomic control and for multiple testing using the more conservative Bonferroni correction. Although we found a number of SNPs that were associated with Ob1, they varied widely in their functions and could not be correlated with any known association with outbreak transmission. Given that Ob2 occurred in the MSM community, we anticipated that there would be one or more identifiable genetic components to explain the focal epidemiology. Unfortunately, the GWAS analysis did not provide data beyond what has been found in other studies; that genomic distinctions between carriage, non-invasive, and invasive isolates of *N. meningitidis* are not easily determined by gene content and may be polygenic [44].

We used a transmission network analysis to evaluate how *N. meningitidis* moved among NYC communities during and between two outbreaks. Isolates from different specimen sources in the same patient were more similar to each other than other isolates in the analysis, demonstrating that within host evolution or variation is not a complicating factor in using the phylogenetics to determine transmission events [45].

Network analysis could assist in resource allocation, such as expanded chemoprophylaxis or vaccination that may abate or interrupt future outbreaks. This methodology can be particularly effective with granular isolate location data, as that is more likely to produce an accurate representation of the movement of bacterial lineages across a population in an area. For example, this concept is demonstrated by the isolate from the Bay Shore, NY patient, which connects to the rest of the locations in different ways in the borough level analysis compared to the neighborhood level analysis. Both the Brooklyn MenC outbreaks began with cases outside of Brooklyn (Ob1: S00-1200 and Ob2: S00-1224) and serve as additional examples (2,3). Whether circulating *Nm* strains, and the individual immunity they provoke in carriers, are localized to neighborhoods or boroughs is not known. It is conceivable that the routine use of network analyses could identify a borough level jump of an *Nm* strain representing an elevated risk of initiating an outbreak.

Brooklyn was identified as an important hub for all *N. meningitidis* infections over the 11 years examined and in particular for the spread of Ob1. There were two transmission events (MN40 to BK37 and MN40 to BK78, Figure 3) within the neighborhood betweenness analysis that were part of the Ob1 outbreak based on epidemiological data but are disconnected from the rest of the outbreak. This is in part because the epidemiological categorization of which isolates are part of an outbreak is imperfect as discussed above. It is also possible for a number of reasons that we are missing isolates that were part of both outbreaks. Carriers of *N. meningitidis* who do not develop meningococcal disease serve as intermediaries between clinical cases, however, they do not get cultured and links in the transmission chain go undiscovered. Similarly, meningococcal cases who are not NYC residents but may work or travel within the five boroughs, are often not reported to DOHMH. One neighborhood in Manhattan was identified as a hub for to the spread of Ob2. Evidence suggests that the Ob2 strain was transmitted to Europe and caused outbreaks in Germany and France [31]. The disconnection in our betweenness network emphasizes the difficulty with performing these types of analyses as a single city. The availability of worldwide transportation allows individuals to move from country to country potentially spreading disease. Perhaps there is no greater recent example than the 2014-5 Ebola virus outbreak in West Africa that spread to Europe and the United States. This only serves to emphasize the importance of countries and institutions sharing data in real-time to obtain the most accurate characterization of global disease movement.

Two *N. meningitidis* outbreaks occurred in NYC in the 21^st^ century and both were centered in Brooklyn. Whole genome sequencing of *N. meningitidis* isolates and betweenness analyses were used to characterize the relationship among cases and illuminate transmission. These tools add an important dimension to our understanding of outbreaks of IMD and have the potential to inform prevention efforts by identifying high transmission areas. One such approach might be to afford two or more IMD cases occurring in space and time in a Brooklyn neighborhood closer scrutiny to uncover links not immediately detected through routine investigations. Additionally, *N. meningitidis* carriage studies in MSM have either been completed (New York City) or are underway (Los Angeles) and will add to our knowledge of the evolution of this serial killer pathogen.

## Acknowledgements

We thank Mike Antwi and Marie Dorsinville who assisted in data collection. We also thank Marcelle Layton, Demetre Daskalakis, and James Hadler for their critical review of the manuscript. This publication made use of the PubMLST website (https://pubmlst.org/) developed by Keith Jolley (Jolley & Maiden 2010, BMC Bioinformatics, 11:595) and sited at the University of Oxford. The development of that website was funded by the Wellcome Trust.

---

**Supplemental Table 1.** New York City Neighborhood Tabulation Areas (NTA) and other Geographic Locations 647 Used in the Betweenness Analysis

| Designation Used<br>In Figures 2 & 3 | Neighborhood/Location<br>Name | New York City<br>Borough |
| --- | --- | --- |
| Bay Shore, NY | Bay Shore, NY | Bay Shore, NY |
| BK25 | Homecrest | Brooklyn |
| BK26 | Gravesend | Brooklyn |
| BK29 | Bensonhurst East | Brooklyn |
| BK31 | Bay Ridge | Brooklyn |
| BK32 | Sunset Park West | Brooklyn |
| BK33 | Carroll Gardens, Columbia Street, Red Hook | Brooklyn |
| BK35 | Stuyvesant Heights | Brooklyn |
| BK37 | Park Slope, Gowanus | Brooklyn |
| BK38 | DUMBO, Vinegar Hill, Downtown Brooklyn, Boerum Hill | Brooklyn |
| BK42 | Flatbush | Brooklyn |
| BK45 | Georgetown, Marine Park, Bergen Beach, Mill Basin | Brooklyn |
| BK61 | Crown Heights North | Brooklyn |
| BK69 | Clinton Hill | Brooklyn |
| BK75 | Bedford | Brooklyn |
| BK78 | Bushwick South | Brooklyn |
| BK81 | Brownsville | Brooklyn |
| BK83 | Cypress Hills, City Line | Brooklyn |
| BK88 | Borough Park | Brooklyn |
| BK93 | Starrett City | Brooklyn |
| BX01 | Claremont, Bathgate | Bronx |
| BX06 | Belmont | Bronx |
| BX13 | Co-Op City | Bronx |
| BX14 | East Concourse, Concourse Village | Bronx |
| BX17 | East Tremont | Bronx |
| BX26 | Highbridge | Bronx |
| BX28 | Van Cortlandt Village | Bronx |
| BX29 | Spuyten Duyvil, Kingsbridge | Bronx |
| BX35 | Morrisania, Melrose | Bronx |
| BX39 | Mott Haven, Port Morris | Bronx |
| BX40 | Fordham South | Bronx |
| BX75 | Crotona Park East | Bronx |
| BX99 | park, cemetery, etc, Bronx | Bronx |
| Central Islip, NY | Central Islip, NY | Central Islip, NY |
| Elizabeth, NJ | Elizabeth, NJ | Elizabeth, NJ |
| Huntington Station, NY | Huntington Station, NY | Huntington Station, NY |
| London, England | London, England | London, England |
| MN04 | Hamilton Heights | Manhattan |
| MN06 | Manhattanville | Manhattan |
| MN12 | Upper West Side | Manhattan |
| MN13 | Hudson Yards, Chelsea, Flatiron, Union Square | Manhattan |
| MN15 | Clinton | Manhattan |
| MN17 | Midtown, Midtown South | Manhattan |
| MN19 | Turtle Bay, East Midtown | Manhattan |
| MN20 | Murray Hill, Kips Bay | Manhattan |
| MN22 | East Village | Manhattan |
| MN34 | East Harlem North | Manhattan |
| MN36 | Washington Heights South | Manhattan |
| MN40 | Upper East Side, Carnegie Hill | Manhattan |
| Mount Croghan, SC | Mount Croghan, SC | Mount Croghan, SC |
| Paris, France | Paris, France | Paris, France |
| QN17 | Forest Hills | Queens |
| QN45 | Douglas Manor, Douglaston, Little Neck | Queens |
| QN61 | Jamaica | Queens |
| QN66 | Laurelton | Queens |
| QN71 | Old Astoria | Queens |
| QN76 | Baisley Park | Queens |
| SI35 | New Brighton, Silver Lake | Staten Island |
| Southold, NY | Southold, NY | Southold, NY |
| Yonkers, NY | Yonkers, NY | Yonkers, NY |

**Supplemental Table 2.**
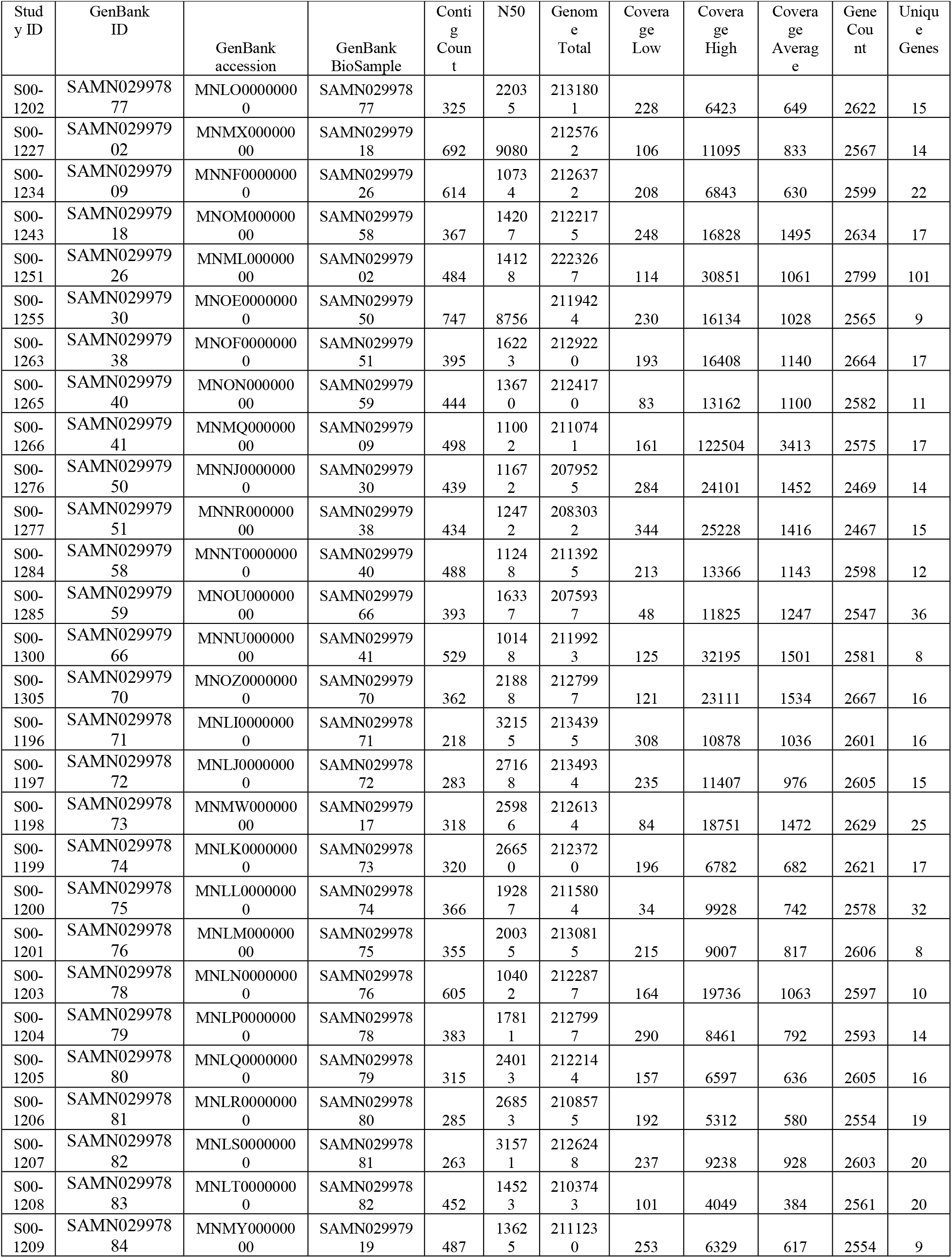
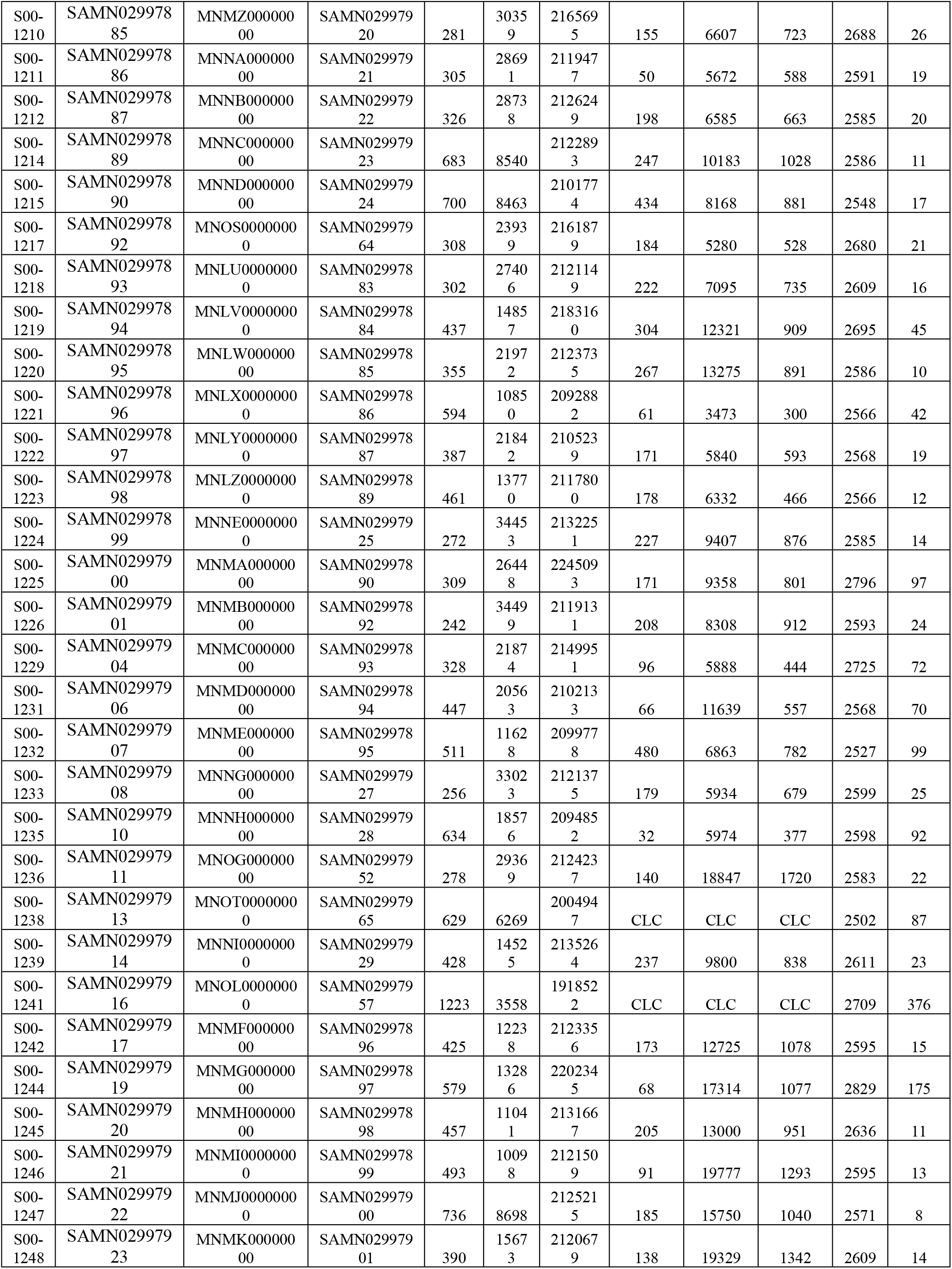
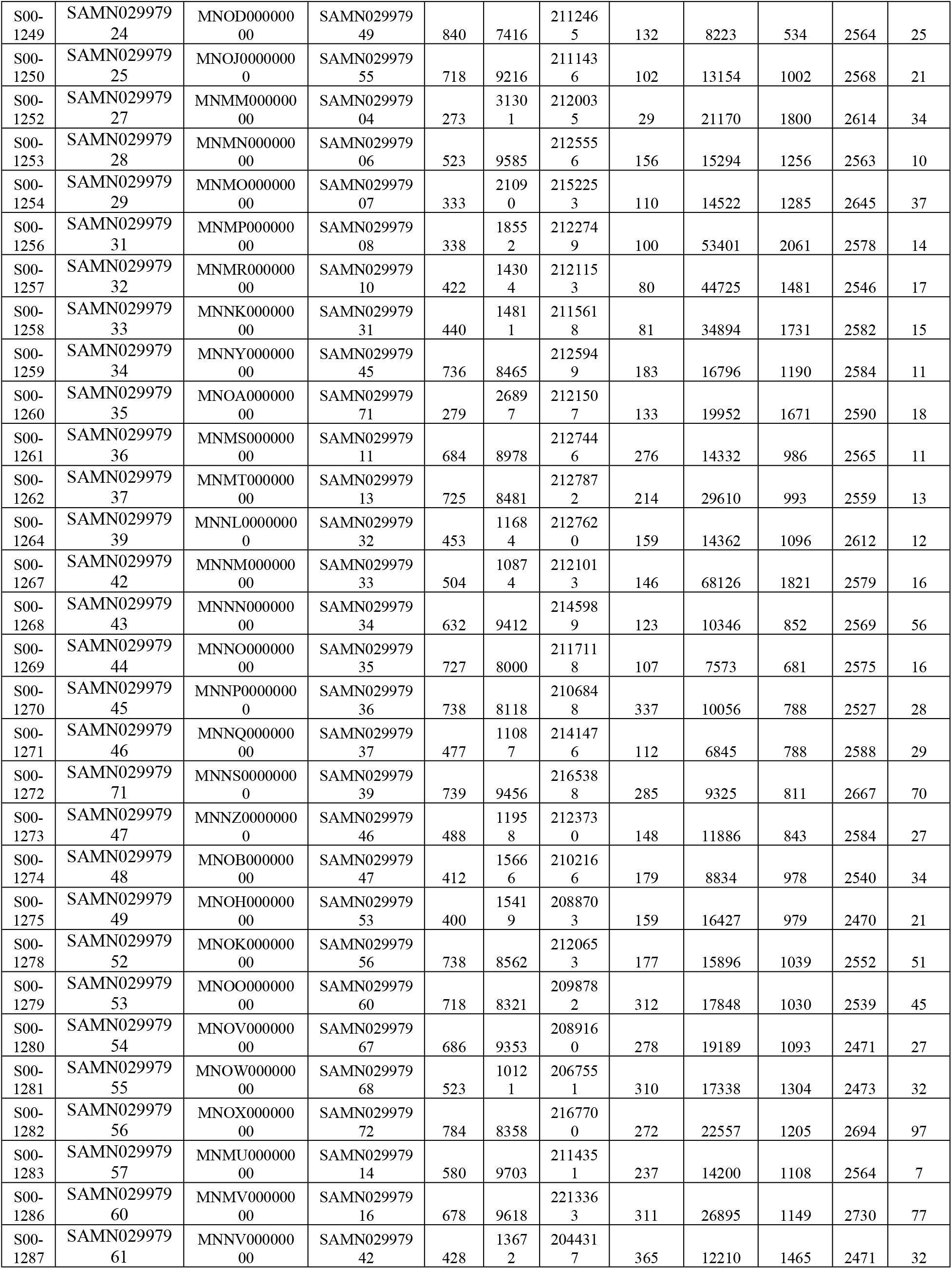
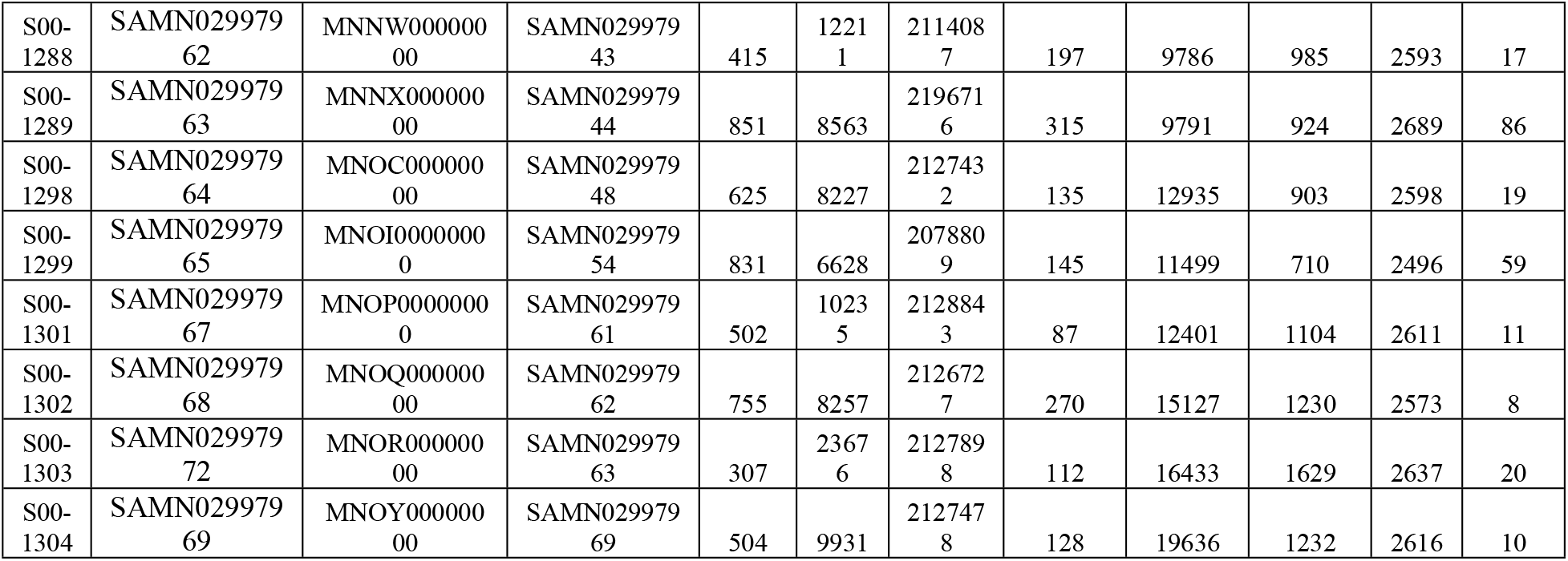
Genomic Metrics of 96 Sequenced *N. meningitidis Isolates*

